# ScisorWiz: Visualizing Differential Isoform Expression in Single-Cell Long-Read Data

**DOI:** 10.1101/2022.04.14.488347

**Authors:** Alexander N. Stein, Anoushka Joglekar, Chi-Lam Poon, Hagen U. Tilgner

## Abstract

RNA isoforms contribute to the diverse functionality of the proteins they encode within the cell. Visualizing how isoform expression differs across cell types and brain regions can inform our understanding of disease and gain or loss of functionality caused by alternative splicing with potential negative impacts. However, the extent to which this occurs in specific cell types and brain regions is largely unknown. This is the kind of information that ScisorWiz plots can provide in an informative and easily communicable manner. ScisorWiz affords its user the opportunity to visualize specific genes across any number of cell types, and provides various sorting options for the user to gain different ways to understand their data. ScisorWiz provides a clear picture of differential isoform expression through various clustering methods and highlights features such as alternative exons and Single Nucleotide Variants (SNVs). Tools like ScisorWiz are key for interpreting single-cell isoform sequencing data. This tool applies to any single-cell long-read RNA sequencing data in any cell type, tissue, or species.

## Introduction

Differential isoform expression between cell types and across conditions plays a major role in the diversification of the proteome (1) and functionality of transcripts in the cell (2). Long-read sequencing has become widely used to address this problem (3–11), and with applications to single-cell isoform sequencing studies (12–16). These approaches have been reviewed in Hardwick et al, 2019 (17). Such data require informative visualizations for single genes, so that the impact of alternative exons, exon combinations, as well as those of transcription start site (TSS) and PolyA sites can be easily appreciated. Here, we present ScisorWiz, a streamlined tool to visualize isoform expression differences across single-cell clusters in an informative and easily-communicable manner. ScisorWiz achieves this with an easy, fast, and reliable method of visualizing differential isoform expression data across multiple clusters and is executable from the command line with the R language (18).

## Usage

ScisorWiz visualizes pre-processed single-cell long-read RNA sequencing data. For a user-specified gene, reads for any number of cell types can be visualized and are clustered by chain of introns (the ordered list of a read’s introns), TSS, and/or PolyA site for each cell type. We have used such plots in our long-read (4, 6, 19) and single-cell long-read publications (12–14). However, customizing such a plot for publication standards includes read mapping, shrinking of introns and recalculation of coordinates, calculation of alternative exons, adjusting plot area depending on number of reads and cell types, as well as plotting Single-Nucleotide Variants (SNVs), insertions, and deletions. This process was previously not automated and was only intended to be used for publication purposes. Now, ScisorWiz does this with a single command in R, allowing for many user-specified options including exploratory, interactive outputs and multiple ways to sort isoforms within each cell type: namely by intron-chain, TSS, PolyA site, as well as all three combined.

ScisorWiz can be run on output generated by scisorseqr (13) or a similarly formatted dataset, which, in turn, can be based on diverse mappers including STAR (20) and minimap2 (21). The first approach uses GFF-files for mappings and read-to-gene assignment files that are generated automatically by scisorseqr. However, the user is free to generate these standardized files by other means. The second method uses more specific files that are intrinsic to scisorseqr - the file in question already contains an assigned gene, TSS, PolyA sites, and the intron- and exon-mappings for each read. Thus, this gene plotting library communicates intimately with scisorseqr. Additionally, through the MismatchFinder function, the data set in question can be compared against the reference genome to determine the locations of SNVs, nucleotide insertions, and deletions to be visualized in the plot.

## Approach

To visualize exons separated by up to ~100-fold larger introns, each purely intronic region is shrunk to 100 bases, while sequences that have annotated or novel exons are displayed with their real size. A drawback of this approach is that short introns (<1 kb) that are fully retained in a long read will be drawn to scale. However, very large introns (»10 kb), for which long reads are unlikely to represent the retained form will be shrunk to 100 bases. By default, the package clusters reads according to intron chains. Reads with identical intron chains are thus displayed together to form exonic blocks. Alternatively, clustering can take into account any combination of TSS, PolyA sites, and intron chains when using scisorseqr-generated files as input. In this situation, only reads with an assigned TSS and/or PolyA site are plotted.

ScisorWiz provides a clear picture of differential isoform expression of genes in any dataset by clustering reads. This reveals differential patterns more clearly, such as alternate exon expression across and within cell types.

## Output

ScisorWiz’s output visualizes isoforms read-by-read for any number of cell types for any user-specified gene. Figure 1 shows *Snap25* gene isoforms across six cell types. Colored boxes are exons per read. For each cell type, reads are ordered by intron chain. Orange exons indicate alternatively spliced exons, defined as being included in at least 5% and at most 95% of overlapping reads taken from the entire data set irrespective of cell type - this range is also user-specified. Consistent with previous observations (13, 22), we find that two neighboring alternative exons in *Snap25* are mutually exclusive. Importantly, we observe this mutual exclusivity to be present in multiple cell types. For higher error-rates such as currently in Oxford Nanopore, 20% and 80% cutoffs provide a clearer picture of alternative exons. There are multicolored dots among the cell types representing the locations of SNVs, insertions, and deletions. By default, only SNVs, insertions, and deletions present in at least 5% and at most 95% of overlapping reads are highlighted in order to avoid plotting random sequencing errors. However, these cutoffs can be adjusted as options by the user allowing the visualization of every single nucleotide disagreeing with the reference genome, should this be of interest. This course of action may be useful in low error-rate sequencing such as Pacific Biosciences (23). Similarly, any mismatches present within the first or last 20 bases of an alignment are not shown in order to avoid alignment artifacts at alignment ends. The bottom section is the GENCODE annotation covered by long reads. ScisorWiz also generates a file for all single-cell long reads that can be uploaded and inspected on the UCSC Genome Browser (24).

**Fig. 1.**
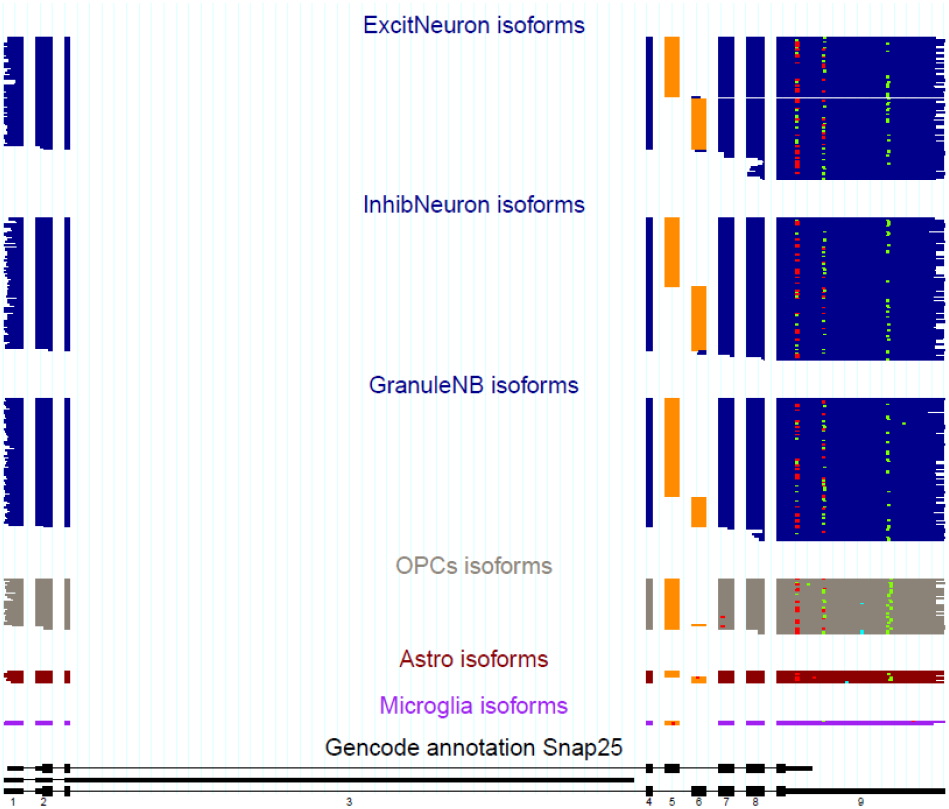
The isoforms of the *Snap25* gene present in each read of a specific cell type are displayed one above the other to form a consistent picture of the gene expression of each cell type. The orange-colored exon represents an exon which is considered alternative as a result of a Ψ value of 5% to 95% inclusion irrespective of cell type. The multicolored dots on the plot represent SNVs (blue), nucleotide insertions (green), and nucleotide deletions (red). All SNVs, insertions, and deletions included are in at least 5% and at most 95% of overlapping reads. The reads at the bottom (black) represent the part of the GENCODE annotation for Snap25.

## Code Availability

Source code is available at github.com/ans4013/ScisorWiz. No new data was generated for this publication. Example data has been included in the package.

## Acknowledgements

We thank the Weill Cornell Medicine Scientific Computing Unit (SCU) for use of their computational resources. This work is supported by NIGMS grant 1R01GM135247-01.

## Notes

https://github.com/ans4013/ScisorWiz

### Competing Interest Statement

The authors have declared no competing interest.

